# Molecular Identification and Characterization of Mobatviruses (*Hantaviridae*) in Lao PDR

**DOI:** 10.64898/2026.04.06.713848

**Authors:** Chittaphone Vanhnollat, Kristina Dimitrova, Longthor Vachouaxiong, Jonathan Audet, Somphavanh Somlor, Sarah J. Medina, Phaithong Bounmany, Khaithong Lakeomany, Vaekey Vungkyly, Gary Wong, Vilakhan Xayaphet, Phetphoumin Paphaphanh, Watthana Theppangna, Bounsavane Douangboubpha, Khamsing Vongphayloth, David Safronetz, Philippe Buchy

## Abstract

Hantavirids, specifically the members within the genus *Orthohantavirus*, represent a significant global public health threat, with bat-associated lineages challenging traditional reservoir paradigms. To investigate the genetic diversity of hantavirids in Southeast Asia, we conducted an expanded surveillance program in Lao PDR from May 2023 to October 2025 in bat populations and wild animals from local wet markets. Using molecular screening and deep sequencing to characterize hantavirids from bat populations and wild animals from local wet markets, we identified 20 positive samples across four bat species, recovering coding-complete genomes for multiple novel variants. Phylogenetic analysis confirmed that these viruses form a monophyletic group within *Mobatvirus*, resolving into two major subclades. The first subclade clustered with Quezon and Robina viruses found in fruit-eating bats. The second subclade further split into two lineages corresponding to Ðakrông and Xuân Sơn viruses, which are associated with trident and leaf-nosed bats, respectively. Despite the strong host specificity observed, the detection of these viruses in a wet market, a critical interface for human-wildlife contact, indicates a potential zoonotic risk. These findings significantly expand the known diversity of mobatviruses in Laos and highlight the urgent need for serological surveillance in at-risk human populations to assess the potential for spillover.

## Introduction

Hantavirids, members of the family *Hantaviridae* (order *Bunyavirales*), are enveloped, negative-sense, tripartite, single-stranded RNA viruses that can pose a substantial threat to global public health [1–3]. While historically associated with rodent reservoirs, the discovery of bat-associated lineages has challenged this paradigm and highlights bats as potential ancestral hosts, given their ecological diversity, long lifespans, and ability to sustain viruses asymptomatically [4,5]. These bat-borne hantaviruses, taxonomically assigned to the genera *Loanvirus* (Lóngquán virus and Brno virus) and *Mobatvirus* (Ðakrông virus [DKGV], Láibīn virus, Lena virus, Nova virus, Quezon virus, Robina virus and Xuân Sơn virus) [1,6–12], were genetically divergent from their rodent-borne counterparts and have been detected across Asia, Europe, Africa, and the Americas, spanning multiple families in both major chiropteran suborders, Yinpterochiroptera and Yangochiroptera [4,10,13]. Despite this broad distribution, their zoonotic potential of these viruses remains poorly understood, study however, serological data provides indirect evidence of possible human exposure [14].

Structurally, hantavirids are pleomorphic virions, typically 80-160 nm in diameter, containing a segmented genome, including small (S), medium (M), and large (L) segments. The viral RNA (vRNA) of each segment consists of an open reading frame (ORF) and a non-coding region (NCR) [15]. The S segment encodes the nucleocapsid (N) protein, which encapsidates the vRNA to form ribonucleoprotein complexes [16–19], while the M segment encodes the glycoprotein precursor (GPC). Cleavage of GPC yields the mature surface glycoproteins Gn and Gc, which mediate viral attachment and membrane fusion [20]. The L segment encodes the viral polymerase, a multifunctional enzyme essential for both genome replication and cap-snatching from host mRNAs [21–24].

Southeast Asia (SEA) has emerged as a region of increasing importance for hantavirus discovery, with serological surveys indicating human exposure across different countries, including Vietnam [25], Thailand [26], Cambodia [27,28], Malaysia [29], Indonesia [30], and the Philippines [31]. However, the region’s potential for harboring novel hantavirids remained largely unexplored. Our previous work in Laos has confirmed the presence of bat-borne hantaviruses closely related to strains circulating in Vietnam [32], expanding the known geographical range of these viruses. However, a comprehensive understanding of the genetic diversity within these viruses remains incomplete, underscoring the need for further sampling and detailed characterization. To address this gap, we continued and expanded the surveillance program for hantaviruses in Laos. By leveraging high-throughput sequencing technologies, we conducted an in-depth genetic analysis of the detected hantaviruses, uncovering their diversity, genomic characteristics, and evolutionary relationships within the context of bat-borne hantaviruses.

## Results

### Sample screening and diversity of bat-borne hantaviruses

Following the initial detection of Lao bat-borne hantaviruses (LBHVs) [32], we expanded our surveillance to assess the prevalence and diversity of hantaviruses at 3 locations in Laos (Fig. 1A). We also incorporated samples collected from wild animals traded in the local wet markets. A total of 1,020 bats were live-captured, and 591 wild animals were surveyed in the market. These animals were classified into 34 genera and 59 species (S1 Fig). From these animals, 821 tissue, 357 intestine, 1,586 anal and 1,589 saliva swab samples were screened for hantaviruses (S2 Fig). Of the total samples tested, 20 samples were positive. Specifically, 17 positives were from tissue samples, while only 2 and 1 were from saliva and anal swab samples, respectively (Fig. 1B). The positive samples originated from 4 different bat species, including 14 from *Aselliscus stoliczkanus* (Stoliczka’s trident bat), 3 from *Hipposideros gentilis* (Andersen’s leaf-nosed bat), 2 from *Rousettus amplexicaudatus* (Geoffroy’s Rousette), and 1 from *Macroglossus sobrinus* (Long-tongued fruit bat). The majority of positive samples (18/20) were from live-captured bats. Notably, 2 samples were collected from a bat found within the local wet market (Fig 1B).

**Figure 1.**
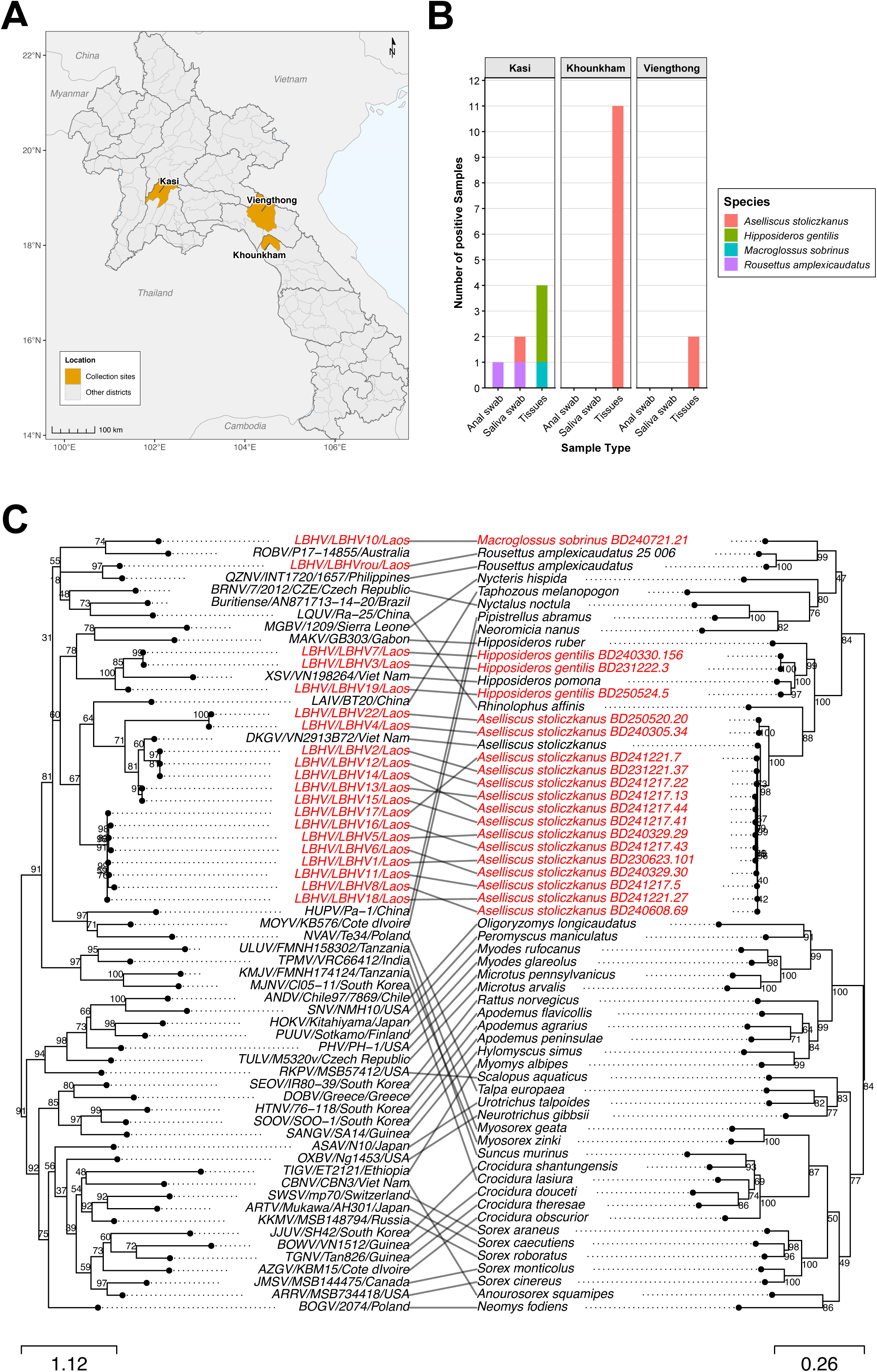
Geographic sampling and host-virus co-phylogeny. (A) Map of collection sites. Detailed coordinates of each location are provided in Methods. (B) Stacked bar charts of PCR-positive samples by location and sample type. Each facet corresponds to a specific site, with colors indicating bat species. (C) Tanglegram of co-phylogeny. The left tree shows the ∼400-bp partial RdRP, and the right shows the ∼1,140-bp bat cytb. Both were reconstructed using maximum-likelihood, and the sequence accession numbers were listed in S3 Table. Lines connect viral sequences to their host counterparts. Scale bars indicate nucleotide substitutions per site.

To assess host-virus evolutionary relationships, we performed co-phylogenetic mapping of bat cytochrome b (*cytb*) and viral partial RdRp gene sequences. The maximum likelihood (ML) tree of the RdRp gene revealed that detected hantaviruses formed three phylogenetic groups sharing clades with other *Mobatvirus* members. The majority of samples clustered with DKGV, the second group was associated with XSV and, the third group grouped with Quezon and Robina viruses (QZNV and ROBV) (Fig 1C). Mapping to the bat *cytb* gene exhibited high host-specificity among the identified mobatviruses, with no evidence of host switching. Specifically, DKGV-related LBHV variants were detected in *Aselliscus stoliczkanus*, XSV-related variants in *Hipposideros gentilis*, and QZNV- and ROBV-related variants in fruit- and nectar-eating bats (*Rousettus amplexicaudatus* and *Macroglossus sobrinus*).

### Genomic description of identified mobatviruses

To maximize the recovery of viral sequences, we applied a host genome depletion prior to nucleic acid extraction and sequencing. From a total of 23 sequencing libraries, including samples from our previous work [32], we successfully obtained coding-complete sequences for the S, M, and L segments of LBHV1, LBHV7, and LBHVrou, as well as the S and M segments of LBHV5 and the M segment of LBHV6. Partial sequences of the S, M, and L segments were obtained for LBHV22, and the M and L segments for LBHV8 (Table 1). For downstream analysis, only intact coding-complete sequences were retained to ensure data integrity and consistency.

**Table 1.** Summary of sampling information and sequencing data analysis.

| Variants | Host species | Date of collection | Province | District | Village | No. of raw reads | No. of total contigs | No. of clustered reads<sup>a</sup> | No. of contigs<sup>b</sup> | Coding-complete sequences |
| --- | --- | --- | --- | --- | --- | --- | --- | --- | --- | --- |
| **LBHV1** | *Aselliscus stoliczkanus* | 23-Jun-23 | Khammouane | KhounKham | Khoun Ngeun | 134,247,408 | 4,098 | 174 | 4 | S, M and L |
| **LBHV2** | *Aselliscus stoliczkanus* | 21-Dec-23 | Khammouane | Khounkham | Khoun Ngeun | 137,213,014 | 9,725 | 53 | 0 | n.d. |
| **LBHV3** | *Hipposideros gentilis* | 22-Dec-23 | Khammouane | Khounkham | Khoun Ngeun | 158,528,330 | 167,626 | 11 | 0 | n.d. |
| **LBHV4** | *Aselliscus stoliczkanus* | 5-Mar-24 | Vientiane | Kasi | Pha Tor | 166,036,074 | 230 | 33 | 0 | n.d. |
| **LBHV5** | *Aselliscus stoliczkanus* | 29-Mar-24 | Khammouane | KhounKham | Khoun Ngeun | 158,119,386 | 267 | 93 | 2 | S and M |
| **LBHV6** | *Aselliscus stoliczkanus* | 29-Mar-24 | Khammouane | KhounKham | Khoun Ngeun | 141,572,766 | 123 | 69 | 2 | M |
| **LBHV7** | *Hipposideros gentilis* | 30-Mar-24 | Khammouane | KhounKham | Khoun Ngeun | 167,395,328 | 80 | 306 | 3 | S, M and L |
| **LBHV8** | *Aselliscus stoliczkanus* | 8-Jun-24 | Bolikhamxay | Viengthong | Vang Hin | 170,226,350 | 262,467 | 104 | 2 | M and L (both partial) |
| **LBHV9** | *Aselliscus stoliczkanus* | 8-Jun-24 | Bolikhamxay | Viengthong | Vang Hin | 280,021,590 | 461,158 | 31 | 0 | n.d. |
| **LBHV10** | *Macroglossus sobrinus* | 21-Jul-24 | Vientiane | Kasi | Pha Tor | 257,424,850 | 172 | 1 | 0 | n.d. |
| **LBHV11** | *Aselliscus stoliczkanus* | 17-Dec-24 | Khammouane | Khounkham | Khoun Ngeun | 245,951,666 | 83,732 | 4 | 0 | n.d. |
| **LBHV12** | *Aselliscus stoliczkanus* | 17-Dec-24 | Khammouane | Khounkham | Khoun Ngeun | 269,223,550 | 254,068 | 3 | 0 | n.d. |
| **LBHV13** | *Aselliscus stoliczkanus* | 17-Dec-24 | Khammouane | Khounkham | Khoun Ngeun | 257,167,374 | 169,465 | 3 | 0 | n.d. |
| **LBHV14** | *Aselliscus stoliczkanus* | 17-Dec-24 | Khammouane | Khounkham | Khoun Ngeun | 263,351,322 | 117,751 | 4 | 0 | n.d. |
| **LBHV15** | *Aselliscus stoliczkanus* | 17-Dec-24 | Khammouane | Khounkham | Khoun Ngeun | 255,875,146 | 233,499 | 17 | 0 | n.d. |
| **LBHV16** | *Aselliscus stoliczkanus* | 17-Dec-24 | Khammouane | Khounkham | Khoun Ngeun | 259,095,288 | 80,127 | 3 | 0 | n.d. |
| **LBHV17** | *Aselliscus stoliczkanus* | 21-Dec-24 | Khammouane | Khounkham | Khoun Keo | 250,638,486 | 1,433 | 3 | 0 | n.d. |
| **LBHV18** | *Aselliscus stoliczkanus* | 21-Dec-24 | Khammouane | Khounkham | Khoun Keo | 281,384,034 | 11,158 | 1 | 0 | n.d. |
| **LBHV22** | *Aselliscus stoliczkanus* | 20-May-25 | Vientiane | Kasi | Pha Tor | 244,443,182 | 6,677 | 152 | 4 | S, M and L (all partial) |
| **LBHV19** | *Hipposideros gentilis* | 24-May-25 | Vientiane | Kasi | Pha Tor | 265,717,912 | 124,813 | 1 | 0 | n.d. |
| **LBHV20** | *Hipposideros gentilis* | 24-May-25 | Vientiane | Kasi | Pha Tor | 240,133,098 | 90,226 | 18 | 0 | n.d. |
| **LBHV21** | *Hipposideros gentilis* | 24-May-25 | Vientiane | Kasi | Pha Tor | 245,509,988 | 118,159 | 5 | 0 | n.d. |
| **LBHVrou** | *Rousettus amplexicaudatus* | 24-May-25 | Vientiane | Kasi | Poukham Market | 327,316,088 | 35,474 | 518 | 3 | S, M and L |

The table includes the sample collection site and date, raw sequencing output, and the results of baseline analysis. n.d., not determined (no coding-complete sequences).
<sup>a</sup>number of hantavirus-assigned clustered reads (from hecatomb pipeline[37])
<sup>b</sup>number of contigs identified as hantavirus

We subsequently compared the amino acid identity (AAI) and average nucleotide identity (ANI) of the coding-complete sequences of LBHV variants against reference mobatviruses and other hantaviruses. The highest AAI between LBHV variants and reference viruses was observed for the S segment (96.9 - 99.0%), followed by M (95.3 - 98.1%) and L segments (93.9 - 97.9%). Specifically, variant LBHV1, 5, and 6 exhibited the highest similarity to DKGV_VN2913B72, while LBHV7 and LBHVrou showed maximal identity to XSV_VN1982B4 and QZNV_MT1720/1657, respectively. The ANI across all segments of LBHV variants followed a consistent trend, reflecting the same variant-specific patterns observed for AAI (Fig 2A).

**Figure 2.**
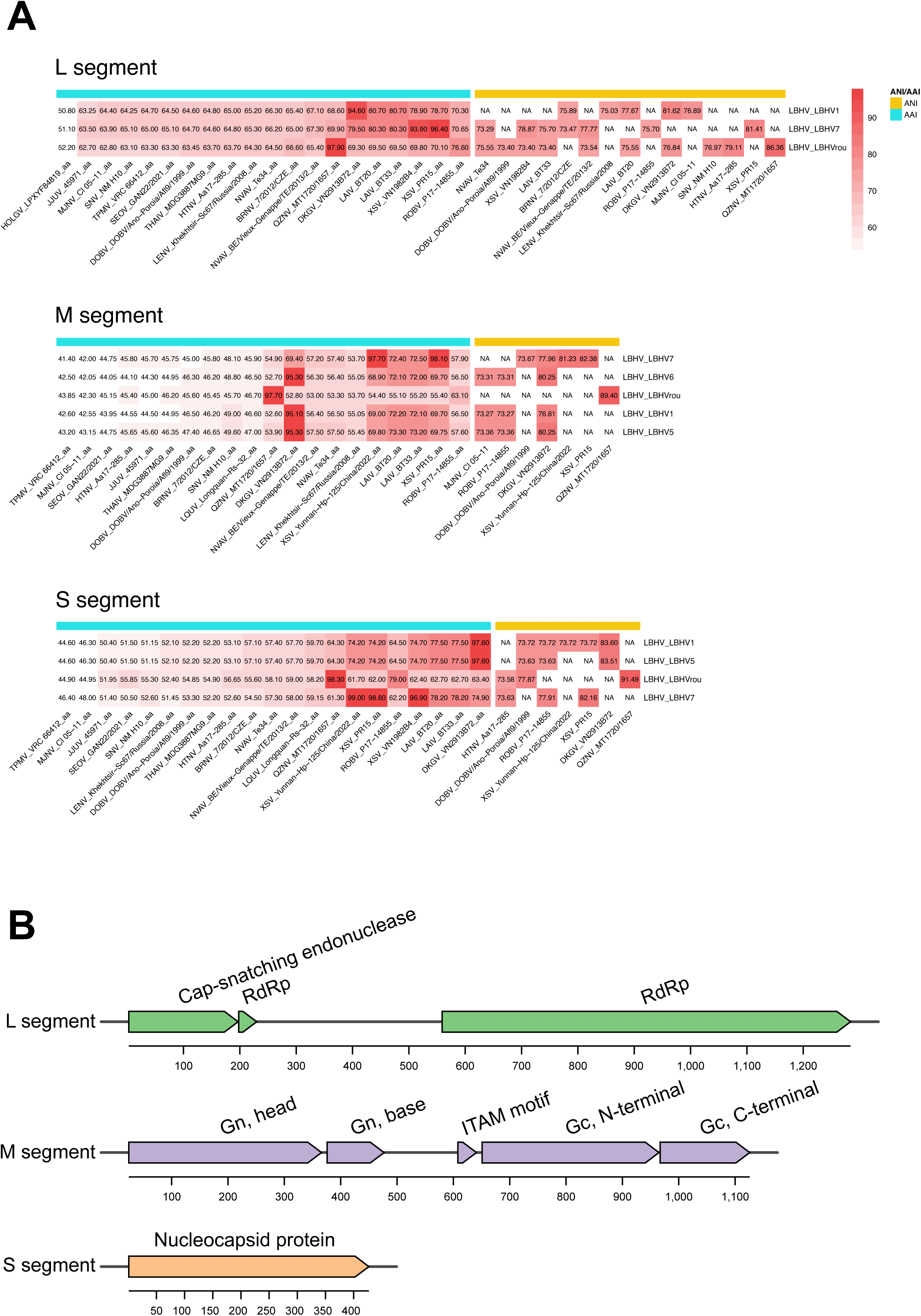
Genome similarity and gene architecture of LBHV genomes. (A) Heatmap of pairwise sequence identity for the S, M, and L segments. Cells marked “NA” indicate values below the fastANI cutoff. (B) Genome organization of LBHV1. The gene track is drawn to scale based on InterProScan coordinates. Detailed tracks for other strains are provided in S1 Table.

### Protein domain prediction and sequence comparative analysis

The protein prediction of LBHV ORF sequences revealed the canonical hantavirus gene organization: the S segment encoded the nucleoprotein (NP), the M segment encoded a GPC that was processed into the N-terminal Gn and C-terminal Gc glycoproteins, and the L segment encoded the RNA-dependent RNA polymerase (RdRp) (Fig 2B). Metrics regarding length, and the start-end residues of both the ORF and protein domain are shown in S1 Table.

We performed a comparative analysis of LBHV sequences with representative hantaviruses from genus *orthohantavirus*, *loanvirus* and *mobatvirus* to compare encoded genes and identify key characteristics, motifs, and residues. The L segment of variants LBHV1, LBHV7, and LBHVrou harbored an endonuclease domain in the N-terminal region (approximately residues 1-200), while the central core contained a canonical RNA-dependent RNA polymerase (RdRp) domain (Fig 3A, S1 Table). The endonuclease domain contained the conserved H…PD…D/ExT…K motif, which was critical for endonuclease function that mediated the cap-snatching activity of hantavirus RdRp [21–24]. The prime-and-realign (PR) loop residues, implicated in de novo initiation of viral mRNA synthesis [51–52], was also observed. The RdRp core contained all conserved polymerase motifs A through G. These motifs followed the typical structural organization of bunyaviral RdRp [21,24]: Motif G (RY residues), Motif F (previously known as Premotif A, contained KxQx₅R…K/Rx₆E residues), motif A (Dx₂KW), motif B (QGx₅SS), motif C (SDD), motif D (KK), and motif E (Ex₂S) (Fig 3A).

**Figure 3.**
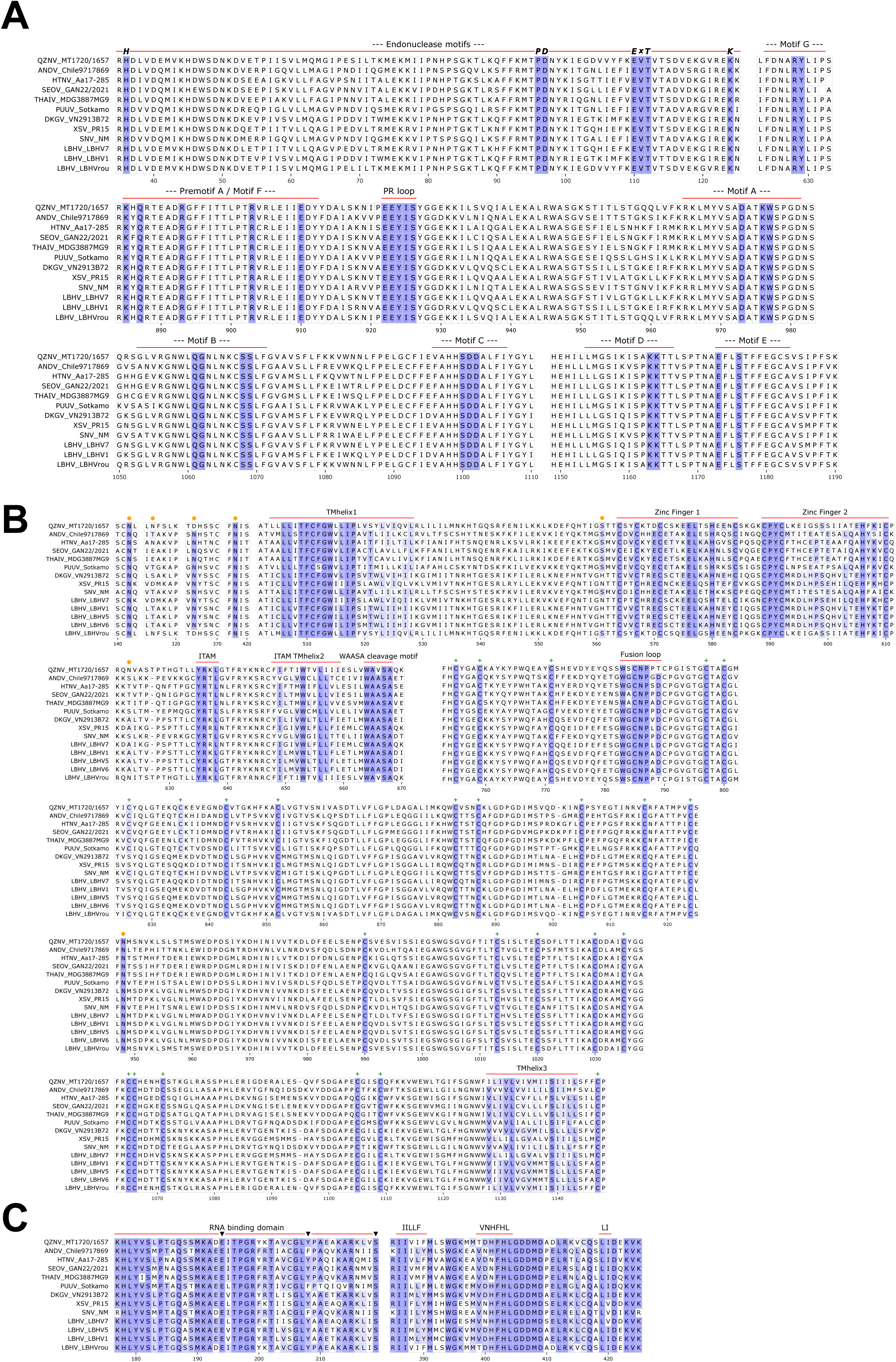
Multiple sequence alignments (MSA) and conserved motifs across segments. MSA of the L (A), M (B), and S (C) segments are shown. The MSA was truncated to highlight specific regions; where conserved motifs and functional regions are highlighted in red lines and annotated above the alignment, while residue conservation (50-100%) was indicated by the intensity of purple. In (B), red-orange dots indicate N-linked glycosylation sites, and green ‘+’ symbols mark cysteine residues. Detailed transmembrane region (TMhelix) is provided in S4 Fig. In (C), black arrows highlight conserved residues.

For M segments, a conserved pentapeptide WAASA was present in variants LBHV1, and LBHV5-7, while WAVSA was only observed in LBHVrou (Fig 3B). The Gn cytoplasmic tail contains a conserved CCHC-type zinc-finger (ZF) domain. Next to the ZF domain, a single immunoreceptor tyrosine-based activation motifs (ITAMs), was observed. The putative fusion loop, Wx_2_Nx_2_D, was present in variant LBHV1, 5-7, whereas Wx_2_Nx_2_T was identified in variant LBHVrou (Fig 3B). Additionally, predicted transmembrane helices were observed in three locations across LBHV variants (S3 Fig). Four conserved asparagine (N)-linked glycosylation sites were observed in all LBHV variants, while an additional site was unique to LBHV7 and XSV, and another was shared by LBHVrou and QZNV, and one unique site was seen in LBHVrou. Furthermore, conserved cysteine residues distributed throughout the Gc ectodomain region were observed (Fig 3B). The predicted secondary structure of LBHV GPC resembled that of other representative hantaviruses (S4 Fig). LBHV GPC comprised of 21.6 – 22.3% of α-helices, 37.7 – 38.2 β-sheet and 39.8 – 40.1 % of random coils (S2 Table, S4 Fig). These characteristics were consistent with glycoproteins of other hantaviruses [7,20,66–69].

The predicted secondary structure of the NP revealed a putative coiled-coil motif consisting of two alpha-helices spanning residues 2–75 in the N-terminal region (S4 Fig). This conserved structure was suggested to contribute to NP-NP trimerization [18,70]. The central domain contained highly conserved residues E192, Y206, and S217 (corresponding to E194, Y208, and S219, in the multiple alignment) (Fig 3C), which were essential for RNA interaction [16,17]. The C-terminal domain (CTD) exhibited more pronounced divergence while preserving helix-loop-helix structure (S4 Fig). Minor substitutions were observed within the three motifs, IILLF, VNHFHL, and LI, that were important for the oligomerization at the CTD [19,71,72]. Specifically, the IIFLY motif was observed in variant LBHV7, IIMLY in LBHV1 and 5, and IIVIF in LBHVrou. Regarding the VNHFHL motif, a substitution of a polar uncharged to a negatively charged residue (N->D) was identified at the second position in LBHV7 (‘TDHFHL’) and LBHVrou (VDHFHL), whereas LBHV1 and 5 retained the conserved VNHFHL sequence. The LI motif was conserved in LBHV1, 5, and LBHVrou, whereas LBHV7 exhibited a conservative substitution to LV (Fig 3C). The overall secondary structure content consisted of 55.2–57.9 % α-helices, 37.6–40.2 % random coils, and 4.3–5.3 % β-strands (S2 Table, S4 Fig).

### Phylogenetic analysis of coding-complete sequences

To refine the higher resolution of phylogenetic relationship of LBHVs with other hantaviruses, we performed the phylogenetic analysis of coding-complete NT and AA sequences of the three segments. Initially, due to limited sequence information of L segment [6,73] and only three LBHV variants (LBHV1, LBHV7 and LBHVrou) for which of the L segment was obtained, the phylogenetic trees were inferred for the concatenated deduced S- and M-segment and individual L segment. All LBHV variants formed a strongly supported monophyletic clade within the genus *Mobatvirus* in the AA-based concatenated SM (bootstrap support = 91%) and L (bootstrap support = 93%) phylogenies (Fig 4A-B). Within this monophyletic clade, the SM AA phylogenetic tree resolved into two major subclades. The first major subclade consisted of LBHVrou, which clustered with ROBV and QZNV. The second major subclade showed additional internal structure and resolved into two minor lineages, with LBHV1, 5 and 6 grouping with DKGV and LBHV7 clustering with XSV. Except for LBHV5 (for which the L segment sequence was unavailable), a comparable and congruent subclade structure was observed in both the L AA and NT phylogenies (Fig 4A-B and S5 Fig), indicating consistent evolutionary relationships at the protein level across these segments. However, in the SM NT tree, the LBHVrou cluster occupied a more basal position (S5 Fig). This discordant placement likely reflected elevated synonymous divergence or rate heterogeneity rather than a distinct evolutionary lineage.

**Figure 4.**
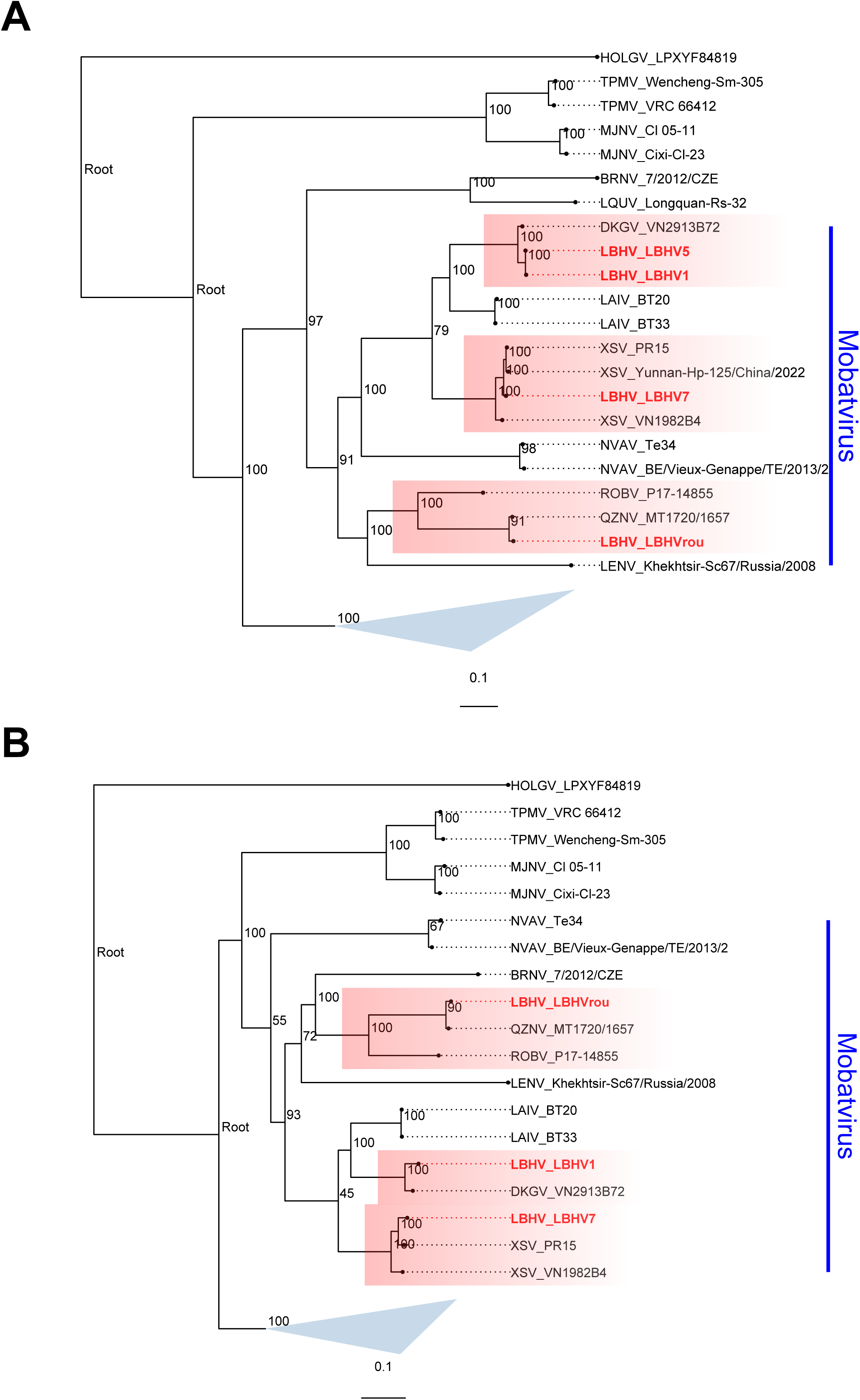
Maximum-likelihood phylogenies of L and concatenated S-M segments. (A) ML tree based on amino-acid sequences of the L segment. (B) ML tree based on concatenated S and M segments. LBHV sequences are highlighted in red, and red rectangles denote major clades. Blue triangle indicates rodent-borne orthohantaviruses. the sequence accession numbers were listed in S3 Table. Scale bars indicate amino-acid substitutions per site. Sequence accession numbers and detailed information of abbreviated tip names are provided in S3 Table.

Next, the NT and AA phylogenies of individual S and M segments were reconstructed to further study the evolution of these segments. The AA and NT phylogenies of M, and NT tree of S segments shared highly similar topological patterns to those of SM AA, and L AA and NT (S5 Fig). However, in the AA phylogeny of S segment, the placement of the LBHVrou cluster was relatively basal compared with other LBHV variants (S5 Fig), reflecting retention of ancestral or distinct protein character states.

### Natural selection

To gain a deeper understanding of the evolutionary dynamics, we performed natural selection analyses using four models to compare LBHV variants with representative hantaviruses. The dN/dS ratio across all three genome segments was extremely low (Table 2), indicating no evidence of pervasive positive selection. MEME identified a limited number of codon sites evolving under episodic positive selection in all three segments. These results suggested that the evolution of these hantaviruses was dominated by strong purifying selection with occasional lineage-specific adaptive events.

**Table 2.** Summary of natural selection analysis.

|  | SLAC |  | FEL |  | FUBAR | MEME |  |
| --- | --- | --- | --- | --- | --- | --- | --- |
| Segments | dN/dS | N<br>(Sites) | dN/dS | N<br>(Sites) | N<br>(Sites) | dN/dS | N (Sites) |
| S | 0.0745 | 0 | 0.0114 | 0 | 0 | 0.0114 | 1 (115) |
| M | 0.0761 | 0 | 0.014 | 0 | 0 | 0.0138 | 6 (10,61,80,860,1009,1050) |
| L | 0.0472 | 0 | 0.0106 | 0 | 0 | 0.0107 | 11<br>(245,329,364,486,796,956,971,1135,1300,1362,1657) |

'N' refers to the number of positively selected amino acids, and 'Sites' indicates the specific amino acid sites under positive selection

## Discussion

### Expansion of *Mobatvirus* diversity and regional host range

Hantaviruses are diverse zoonotic pathogens with a broad host range [4,7–9], yet the diversity of the *Mobatvirus* genus in Southeast Asia remains underexplored. In this study, we identified two additional bat species, *Rousettus amplexicaudatus* and *Macroglossus sobrinus*, that carry mobatviruses in Laos. The detection of positive samples in Vienthong, a district adjacent to Khounkham, where bat-borne hantaviruses were first detected, indicates that the distribution of LBHV may be broader than currently recognized. Together with our previous finding of LBHV in *Aselliscus stoliczkanus*, *Hipposideros gentilis* [32], this study expands the known reservoir diversity and geographical distribution of mobatviruses in this region. In addition, this study further supports our previous observation that recurring detection of highly similar viruses at these sites over time may indicate endemic circulation within these areas, as opposed to sporadic transmission via bat migration [32].

### Phylogenetic placement and host-virus relationship

The phylogenetic analysis of coding-complete sequences provided higher resolution than partial gene analysis. Our data confirms that all identified LBHVs form a monophyletic group within the genus *Mobatvirus* (Fig 4A-B). Notably, the full-genome phylogeny identified two major subclades. The first subclade, containing LBHVrou, clustered with QZNV. While QZNV was previously identified in the Philippines [9], the detection of this cluster in Laos raises questions regarding its origin. It remains questionable whether this represents a recent viral expansion due to bat migration or a long-standing divergence that has been obscured by insufficient sampling. Although *Rousettus amplexicaudatus* is distributed across SEA countries (including Laos and the Philippines) [74], the observed sequence divergence (86-91% at AA level and 97-98% at NT level) (Fig 2) and evidence of purifying selection (Table 2) suggests that the detection of these two phylogenetically related lineages reflects ancient divergence rather than host-mediated dispersal. However, further sampling and in-depth phylogeographic analysis are still required for definitive confirmation.

The second subclade consisted of two internal lineages. First lineage grouped LBHV1, 5, and 6 with DKGV, whereas another lineage grouped LBHV7 with XSV (Fig 4A-B). These clusters are particularly compelling, given the host specificity observed in the region, in which DKGV has been exclusively documented in *Aselliscus stoliczkanus* [7,32] while XSV has been found in *Hipposideros gentilis* (previously known as *Hipposideros pomona*) and *Hipposideros cineraceus* [8,11,31]. This phenomenon is supported in this study by the repeated detection of DKGV-related and XSV-related mobatviruses in bat populations of the same species across two longitudinal sampling sites (Fig 1B-C and Table 1). This finding on the strict host-virus relationship supports the established knowledge that highly similar hantaviruses are primarily circulating in similar host species [5].

### Genomic characteristics and unique features

The genomic organization of the LBHV variants follows the functional features with the other hantaviruses, characterized by a highly conserved L-segment replication machinery, indicated by the presence of canonical endonuclease motif (H…PD…D/ExT…K) and RdRp motifs A-G (Fig 2B, Fig 3A) [21–24]. The M segment analysis shows that LBHV variants use a Class II fusion mechanism identical to other rodent-borne counterparts [20,68,75]. The presence of the highly conserved Wx2Nx2D/T fusion loops in the Gc glycoprotein suggests that pH-dependent membrane fusion remains a strictly preserved trait in LBHV variants (Fig 3B) [7,68,75]. While the typical pentapeptide WAASA is observed in LBHV1 and 5-7, LBHVrou contains a WAVSA variant (Fig 3B). This conservative substitution is unlikely to alter the Gn/Gc cleavage efficiency; rather, it appears to reflect a pattern of host-virus co-divergence, as the WAVSA variant has also been observed in QZNV, which shares the same bat species as LBHVrou (Table 2). Furthermore, the identified LBHV variants lack a complete ITAM (Fig 3B), a common feature of Hantavirus Pulmonary Syndrome (HPS)-causing hantaviruses [69]. The observed single YxxL motif mirrors the genetic profile of Hemorrhagic Fever with Renal Syndrome (HFRS)-associated and non-pathogenic orthohantaviruses. Analysis of the S segment underscores the structural integrity of the NP, which uses a conserved N-terminal coiled-coil for trimerization and essential residues (E192, Y206, S217) for RNA interaction [17]. However, the CTD reveals significant plasticity; specifically, the polar-to-acidic substitutions (N -> D) observed in the V(N/D)HFHL motifs of LBHV7 and LBHVrou may influence RNP assembly or stability [71]. Collectively, although LBHV variants possess typical protein characteristics of hantaviruses, variant-specific variations have been observed. These findings reflect distinct evolutionary pressures or reservoir adaptations. Moreover, further studies for receptor usage and receptor binding sites of the identified bat-borne hantaviruses are imperative to gain a deeper understanding of virus attachment and cell entry, and most importantly, whether they can enter human cells.

### Virus isolation attempts

In addition to the genetic characterization, we attempted isolation using multiple cell lines, including Vero E6, Vero 76, VERO, and the free-tailed bat lung cell line Tb1Lu (CCL-88). Despite six passages, no successful isolation was achieved, as determined by PCR and monitoring for cytopathic effects. This outcome is consistent with previous reports on unsuccessful efforts to isolate bat-borne hantaviruses [12,76]. Considering the high host-specificity of these viruses (Fig 1B-C), future virus-isolation attempts using reservoir host-derived cell lines could be a more effective approach [4,77].

### Zoonotic risks

Given the endemic presence of these viruses in local bat populations and the cultural practice of bushmeat consumption in Laos [78], the detection of mobatviruses in a local wet market, a known hotspots for human-animal interface interactions, may represent a potential zoonotic risk (Fig 1B and Table 1). While the low dN/dS ratios indicate strong purifying selection, suggesting that the viruses are stable and well-adapted to their bat reservoirs rather than rapidly adapting to new hosts (Table 2), the presence of the virus at a high-contact environment warrants close attention and further serological surveillance in humans and market workers to assess the actual zoonotic threat.

## Conclusion

In summary, this study significantly expands the diversity and genomic landscape of *Mobatvirus* in SEA. We identified distinct lineages with unique genomic features. The absence of successful viral isolation underscores the technical challenges inherent in culturing bat-borne hantaviruses. However, the detection of these viruses in a wet market, critical interfaces for human-wildlife contact, highlights the need for continued surveillance to monitor zoonotic risks. Future studies should focus on the functional characterization of these mobatviruses and serological screening of human and animal populations to better understand the public health implications of these emerging bat-borne pathogens.

## Materials and Methods

### Sample collection and PCR screening

The approval to conduct the field sample collection was authorized by the animal health authorities of the Department of Livestock and Fisheries (DLF), Ministry of Agriculture and Forestry (MAF), Lao PDR. The sampling procedure has been previously described [32] with slight modifications. In brief, field sample collections were conducted at 3 sites: Khounkham district in Khammouane Province (18.16N, 104.47E), Viengthong district in Bolikhamxay province (18.30N, 104.50E) and Kasi district in Vientiane Province (19.13N, 102.12E) during May 2023 – October 2025. A study area map was generated using R (v4.5.2) with the ggplot2 and sf packages (Fig. 1A). Administrative boundaries for Laos (provinces and districts) were obtained from Open Development Mekong [33], and neighboring country borders were sourced from Natural Earth (https://github.com/ropensci/rnaturalearth). Geographic coordinates were standardized to the WGS84 CRS (EPSG:4326).

In live-capturing events, bats were captured using 4-bank harp traps and mist nets that were set 1 hour before dusk until dawn. The taxonomy of all captured bats was morphologically identified using key characteristics by a bat taxonomist at the time of collection, as previously described [34]. Saliva and anal swabs, tissue (consisting of pooled heart, lung, liver, spleen, and kidney tissues), and intestine samples were collected. For the local wet market surveys, facilitated by the local authorities, the sampling site was set near the markets. With the permission of the vendors, swab samples were collected from wild animals found in the market. Photos of the sampled animals were captured, and their taxonomy was identified later by taxonomy specialists. All field members participated in bat handling and sample collection, at a minimum, wore eye protection, non-valved N95 respirators, fluid-resistant protective clothing, and double gloves. Protocols for sampling and euthanizing captured animals were performed in accordance with the Canadian Council on Animal Care (CCAC) guidelines: wildlife (https://ccac.ca/Documents/Standards/Guidelines/CCAC_Guidelines-Wildlife.pdf).

For total nucleic acid extraction, tissue samples were homogenized using Omni Bead Ruptor Elite homogenizer at 4 m/s for 30 seconds. Then, the homogenates and swab samples were further extracted for total DNA and RNA using the NucleoSpin 8 virus kit (Macherey Nagel). Reverse transcription was performed using the Maxima H minus first strand cDNA synthesis kit (Thermo Scientific) following manufacturer’s instructions. Samples were tested for the presence of hantavirus by nested PCR [35]. PCR products were subsequently sequenced on both strands by Sanger sequencing with the second-round PCR primers. The obtained sequences were confirmed by homology search using NCBI BLASTn (http://www.ncbi.nlm.nih.gov/BLAST).

### Virus isolation

For virus isolation studies, multiple cell lines were cultured in parallel for comparative analysis, including several Vero E6 strains, Vero 76, VERO cells, and the free-tailed bat lung cell line Tb1Lu (CCL-88). Virus isolation was attempted in the Containment Level 3 laboratories at the National Microbiology Laboratory of the Public Health Agency of Canada, Winnipeg. Each cell line was seeded into 12-well plates one day before the isolation attempt to ensure confluence within 24 hours. Inoculations were performed using a 1% bat tissue homogenate derived from field-collected samples. Following a 1-hour adsorption period at 37 °C, culture medium containing reduced FBS (2%) was added, and plates were incubated at 37 °C with 5% CO₂ for up to 10 days prior to first passage. On day 7 post-inoculation, 140 µL of cell culture supernatant was collected for RNA extraction and PCR analysis. On day 10, cells were collected, pelleted by centrifugation, and harvested. An additional 140 µL of supernatant was aliquoted for RNA extraction and PCR analysis, and the remaining inoculated cell suspension was passaged onto fresh 12-well plates containing the corresponding cell lines. The procedure was carried out through six passages, after which PCR results indicated unsuccessful virus cultivation. Electron microscopy demonstrated cell line-dependent morphological variation; however, consistent with previous hantavirus isolation attempts, no cytopathic effects (CPE) were observed.

### High-throughput sequencing

For high-throughput sequencing (HTS), the homogenized samples tested positive by PCR were subjected to filtration and enzyme cocktail treatment prior to total nucleic acid extraction to remove the eukaryotic/bacterial cells and naked DNA as described in detail elsewhere [36]. Briefly, the homogenates were initially filtered using 0.45 μm PVDF (Milipore). Then, the filtrates were further incubated with 14 U Turbo DNase (Thermo Fisher), 20 U RNase ONE (Promega), and 20 U Benzonase (Sigma) at 37°C for 30 minutes. The total nucleic acid extraction was performed as mentioned above. The sequencing libraries were prepared using the SMARTer Stranded Total RNA-seq kit v3-Pico input mammalian kit (Takara Bio, San Jose, CA, USA) following manufacturer’s instructions. The quantity and size of the libraries were assessed by the Qubit DNA high-sensitivity assay (Invitrogen) and the TapeStation High Sensitivity D1000 assay (Agilent), respectively. The sequencing was performed using NovaSeq 6000 at Macrogen Asia Pacific, Singapore.

### HTS data analysis

Raw reads were bioinformatically processed using the Hecatomb pipeline v1.3.2 [37]. First, adapters and low-quality bases were removed using fastp [38], host genomic sequences were filtered by minimap2 [39], and reads were clustered with MMseqs linclust [40,41]. The clean reads were then annotated using MMseqs2 against viral, multi-kingdom, and polymicrobial amino acid and nucleotide databases. De novo assembly of the pre-processed reads was carried out using Megahit and Flye [42,43]. Finally, the assembled contigs were taxonomically annotated in combination with the information from read-based annotations. The open reading frames (ORFs) of hantavirus-assigned sequences were identified using command-line version of NCBI’s ORFfinder (https://www.ncbi.nlm.nih.gov/orffinder/, accessed on 22 March 2025). For any sequences that had a gap in the middle, primers were designed ad hoc to fill in those gaps. The protein domain of ORFs was identified using InterProScan 5.73-104.0 [44]. The mean coverage of sequences was performed using the mapping-based method integrated in Koverage [45]. To extract cytochrome b (*cytb*) sequences from HTS data, MitoZ v3.6 was initially used [46]. Subsequently, these *cytb* sequences were applied as references for mapping trimmed reads using minimap2 [39], and a consensus sequence was generated with samtools v1.23 [47].

### Sequence similarity and comparative analysis

The similarity of nucleotide and amino acid sequences was assessed using FastANI v1.34 and EzAAI v1.2.3 [48,49], respectively. The secondary structure diagrams were generated from Alphafold3-predicted monomer structure using SSdraw [50,51]. The transmembrane regions of the M segment were estimated by DeepTMHMM v1.0.24 through pybiolib Python package, and the glycosylation sites were identified using the NetNGlyc v1.0 webserver (https://services.healthtech.dtu.dk/) [52,53], respectively.

### Phylogenetic inferences

The multiple sequence alignment was performed using MAFFT v7.525 [54] and the poorly aligned or uninformative sites were trimmed using BMGE (Block Mapping and Gathering with Entropy) v 1.12 [55]. IQ-TREE v2.4.0 was used to infer the maximum likelihood phylogeny with 1000 ultrafast bootstrap replicates [56]. The best-fit model of substitution was identified using the ModelFinder [57]. The tree visualization and annotation were carried out using R package ggtree v3.16.3 [58]. For co-phylogeny reconstruction, trees were inferred using the same method described previously. The tanglegram was then generated using the ‘cophylo()’ function from the phytools R package [59].

### Natural selection analysis

Selection pressures acting on viral coding sequences were assessed using codon-based models implemented in HyPhy [60] and accessed via the Datamonkey web server [61] (https://www.datamonkey.org/). Site-specific selection was evaluated using Single Likelihood Ancestor Counting (SLAC), Fixed Effects Likelihood (FEL), Mixed Effects Model of Evolution (MEME), and Fast Unconstrained Bayesian Approximation (FUBAR) [62–64]. SLAC and FEL were used to detect pervasive selection across lineages, while MEME was employed to identify episodic diversifying selection acting on a subset of branches. FUBAR provided Bayesian estimates of pervasive selection that are robust to model misspecification. Statistical significance was assessed using P < 0.05 for SLAC, FEL, and MEME, and a posterior probability > 0.9 for FUBAR. A ratio of nonsynonymous to synonymous substitutions (dN/dS) was < 1 and interpreted as evidence of purifying selection, whereas dN/dS > 1 indicated positive selection.

## Acknowledgements

We thank Kedkeo Intavong, Souksakhone Viengphouthong, Thep Aksone Chindavong, and Sitsana Keosenhom for their technical support in assisting with sample screening, and Khampok Phithakthep for coordinating and supporting the field missions. We are grateful to the local authorities and communities in Kasi, Khounkham, and Viengthong districts for their cooperation and support throughout this study.

## Data availability

All data are provided within the manuscript and its Supporting Information files. The sequences obtained in this study were submitted to NCBI Genbank with accession numbers: PZ241949-PZ241960.

## Supporting information

**S1 Fig.**
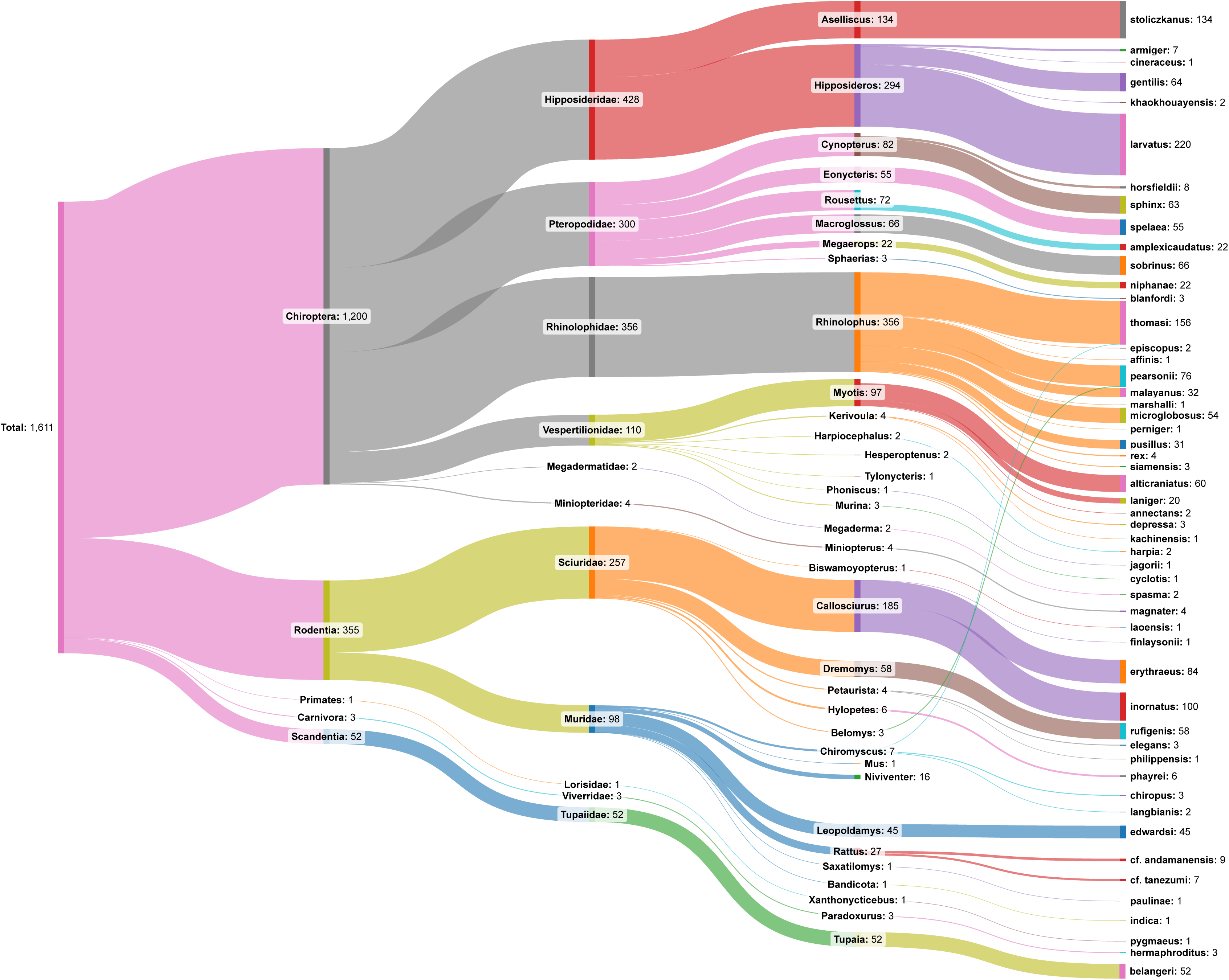
Sankey diagram summarizing number of animals captured and surveyed with taxonomic information. The diagram flows from total samples -> order -> family -> genus -> species.

**S2 Fig.**
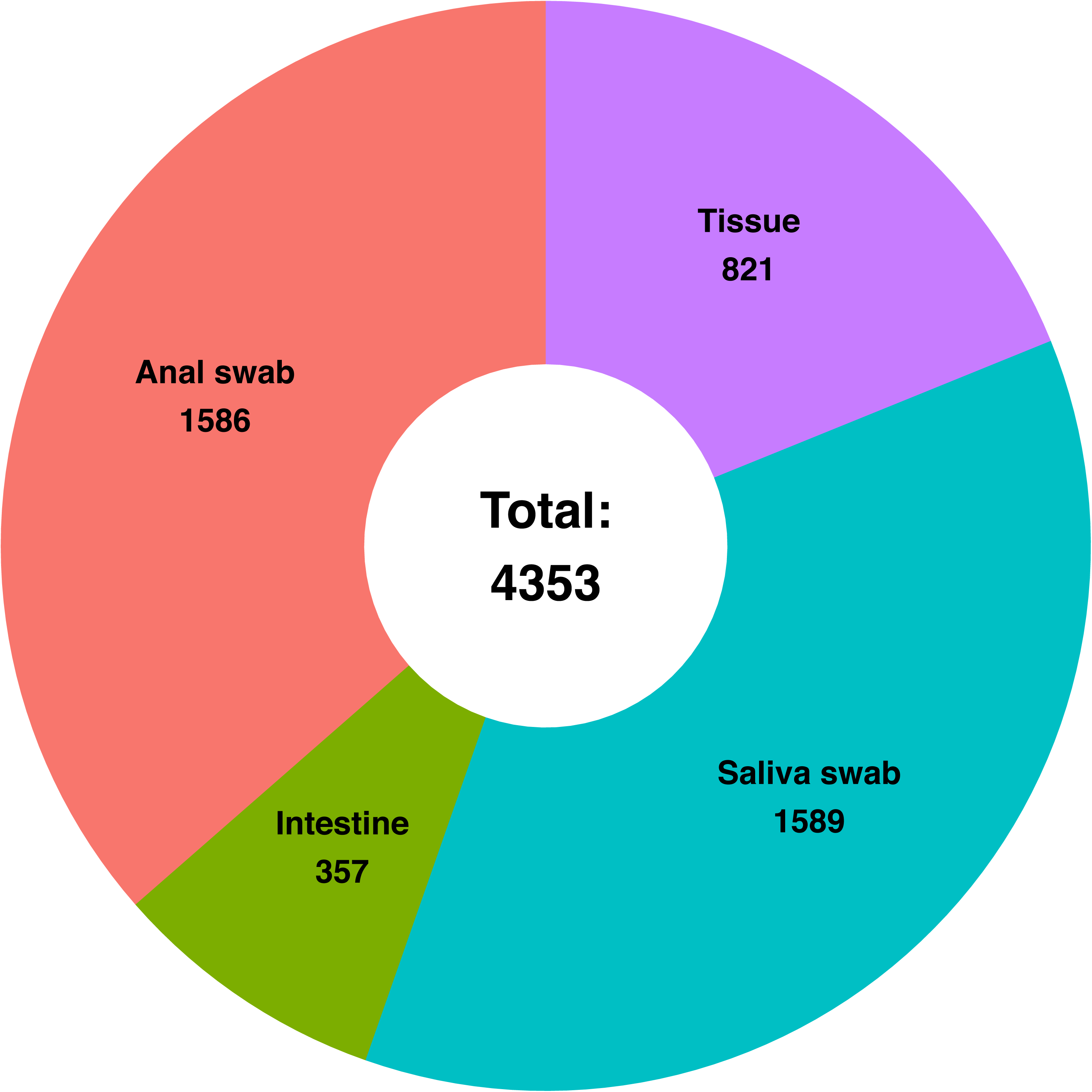
Pie chart showing the composition of the screened samples. Slices represent sample types with counts indicated for each slice.

**S3 Fig.**
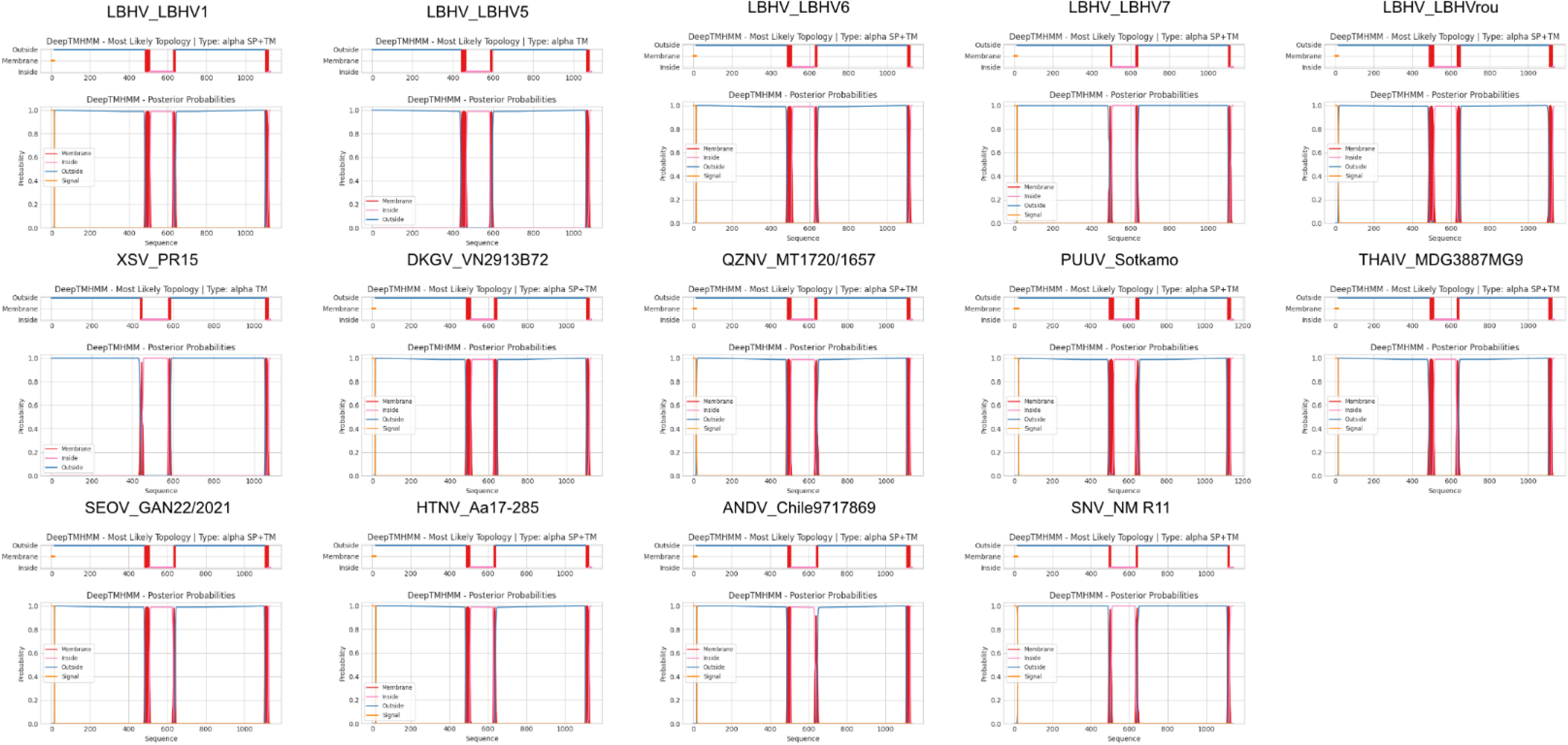
Comparison of predicted transmembrane structure of glycoprotein for LBHV and representative hantavirids. Representative hantavirus sequences used for comparison are listed in S3 Table.

**S4 Fig.**
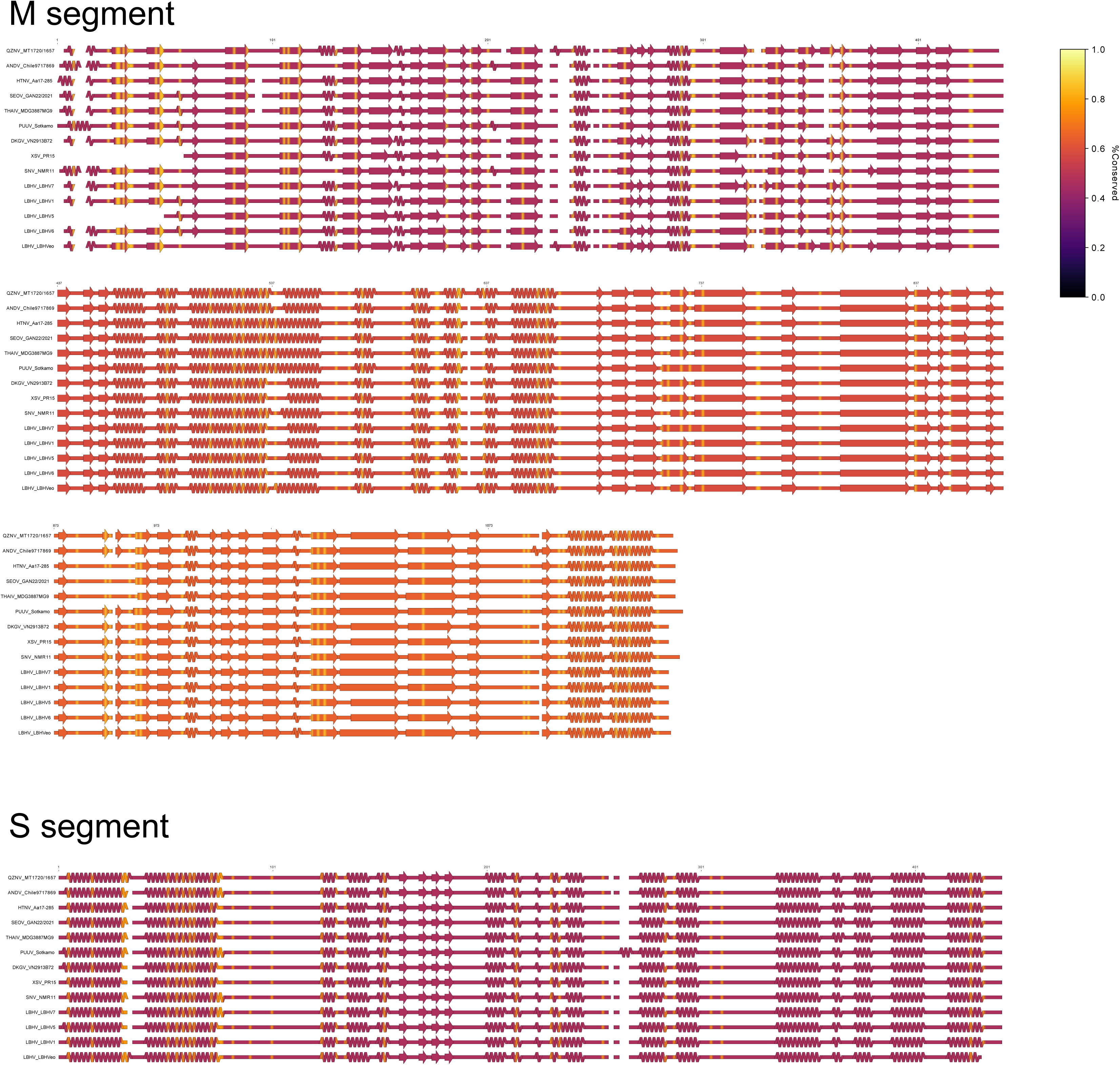
Predicted secondary structure maps for the S and M segment proteins. Secondary structure elements were extracted from AlphaFold3-predicted models (see Methods and S2 Table). Helices, sheets and coils are indicated by standard schematic symbols; conservation (% conserved) was indicated by ssdraw ‘-conservation_score’ parameter.

**S5 Fig.**
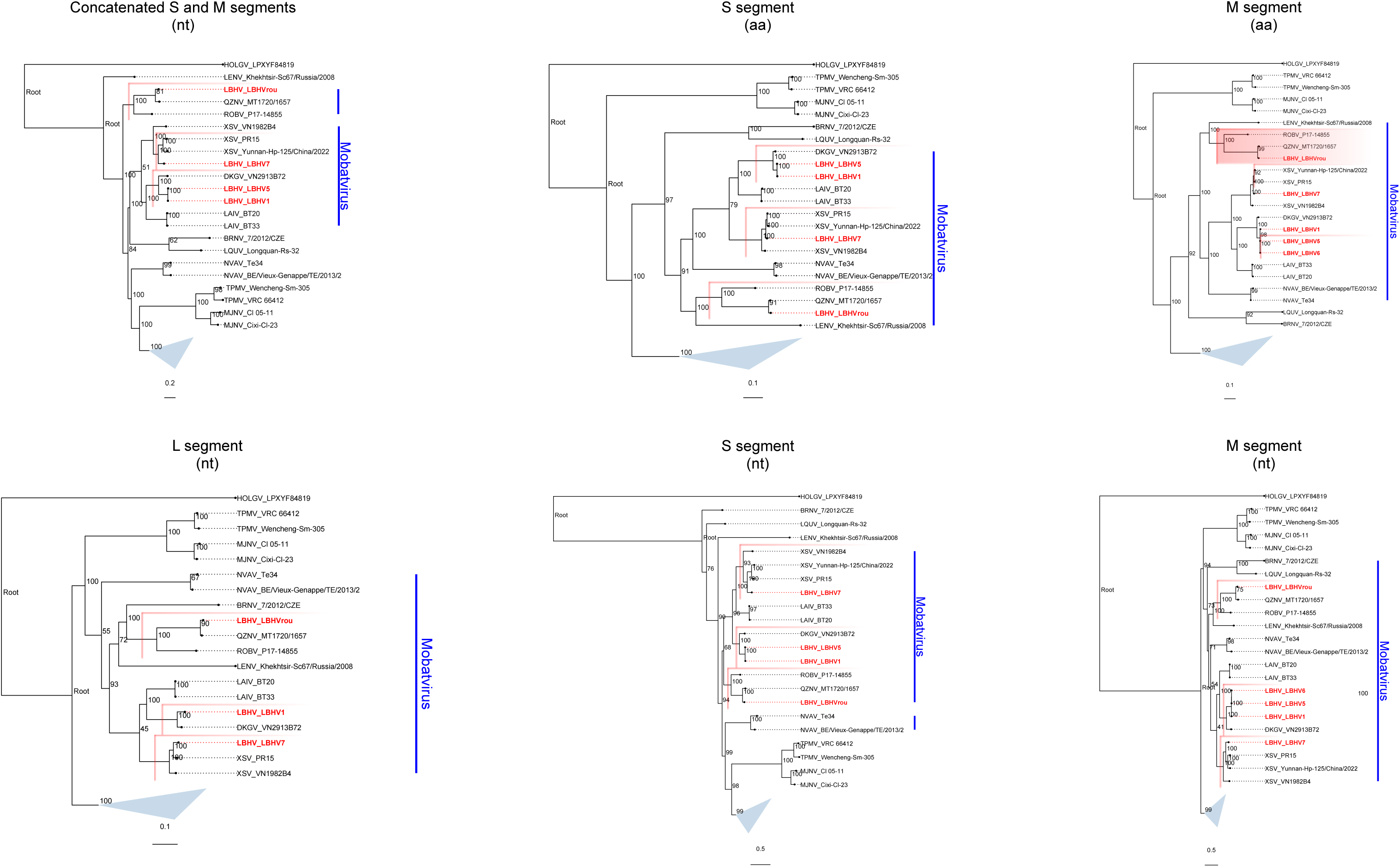
ML trees of both nucleotide (nt) and amino-acid (aa). LBHV sequences are highlighted in red, and red rectangles denote major clades. Blue triangle indicates rodent-borne orthohantaviruses. Scale bars indicate nucleotide substitutions per site for nucleotide trees and amino-acid substitutions per site for amino-acid trees. Sequence accession numbers and detailed information of abbreviated tip names are provided in S3 Table.

S1 Table. Predicted protein domains for LBHV genomes. Table lists each predicted protein, start-end coordinates, matched domain name, and brief functional notes extracted from the InterProScan annotation.

S2 Table. Summary of secondary structure content derived from AlphaFold3-predicted models. The content was extracted using MDAnalysis python library (https://www.mdanalysis.org/).

S3 Table. A list of accession numbers and associated metadata for all sequences used in this study. n.a., not applicable.

